# Do Existing COVID-19 Vaccines Need to Be Updated in 2025?

**DOI:** 10.1101/2025.05.02.651777

**Authors:** Ian A. Mellis, Madeline Wu, Qian Wang, Anthony Bowen, Carmen Gherasim, Riccardo Valdez, Jayesh G. Shah, Lawrence J. Purpura, Michael T. Yin, Aubree Gordon, Yicheng Guo, David D. Ho

## Abstract

COVID-19 vaccines have been updated each year since 2022 to improve protection against evolving SARS-CoV-2 variants. However, it is unclear whether a reformulation will be necessary for 2025. KP.2-based monovalent COVID-19 mRNA vaccines (KP.2 MV) were authorized for use in 2024, and they conferred substantial protection against hospitalizations caused by viral variants that emerged and dominated later, such as KP.3.1.1 and XEC. Today, LP.8.1 and its subvariant LP.8.1.1 have become dominant worldwide, particularly so in North America. Other variants, such as the LF.7 subvariant LF.7.2.1, have emerged with a growth advantage in Asia. To characterize the antigenicity of LP.8.1, LP.8.1.1, LF.7, LF.7.2.1, and another variant under monitoring, MC.10.1, we tested serum samples from 20 individuals who recently received KP.2 MV in neutralization assays against JN.1, KP.2, KP.3, KP.3.1.1, XEC, LP.8.1, LP.8.1.1, LF.7, LF.7.2.1, or MC.10.1 pseudoviruses. Serum neutralizing antibody titers against LP.8.1, LP.8.1.1, LF.7, LF.7.2.1, and MC.10.1 were comparable to those against KP.3.1.1 and XEC, indicating that LP.8.1.1 and other recently dominant subvariants are antigenically similar to their predecessors. Therefore, the currently authorized KP.2 MV may not need to be updated for 2025, if the vaccine manufacturers could demonstrate comparable immunogenicity for KP.2 MV and LP.8.1-based mRNA vaccines and, of course, in the absence of an antigenically divergent SARS-CoV-2 variant emerging.

## Main Text

COVID-19 vaccines have been updated each year since 2022 to improve protection against evolving SARS-CoV-2 variants^1^. However, it is unclear whether a reformulation will be necessary for 2025. KP.2-based monovalent COVID-19 mRNA vaccines (KP.2 MV) were approved in late 2024 in the US, EU, and UK to improve protection against the antigenically distinct viruses in the JN.1 lineage, including KP.2 and KP.3 that were prevalent at the time of strain selection. Indeed, KP.2 MV robustly boosted neutralizing antibody titers against not only the intended target viruses but also ones that emerged later and became dominant, such as KP.3.1.1 and XEC^1^. Recent work showed that KP.2 MV have substantial vaccine efficacy against KP.3.1.1 and XEC, with estimates of up to 75% in protecting against hospitalization in those 65 years of age or older^2^. A progeny virus, LP.8.1, emerged in early 2025 and it and its descendant LP.8.1.1 became dominant worldwide in the ensuing months (Figs. 1A, 1B, S1A). Today, the LP.8.1 sublineage accounts for >70% of new SARS-CoV-2 infections in North America, based on viral sequences deposited (Fig. 1B). LP.8.1 possesses 5 additional spike mutations compared with KP.3.1.1, including R346T and H445R in the receptor-binding domain, F186L and R190S in the N-terminal domain, as well as K1086R in the S2 region (Figs. 1A, S1B). LP.8.1.1 bears an additional K679R mutation near the Furin cleavage site (Fig. S1B).

**Figure 1:**
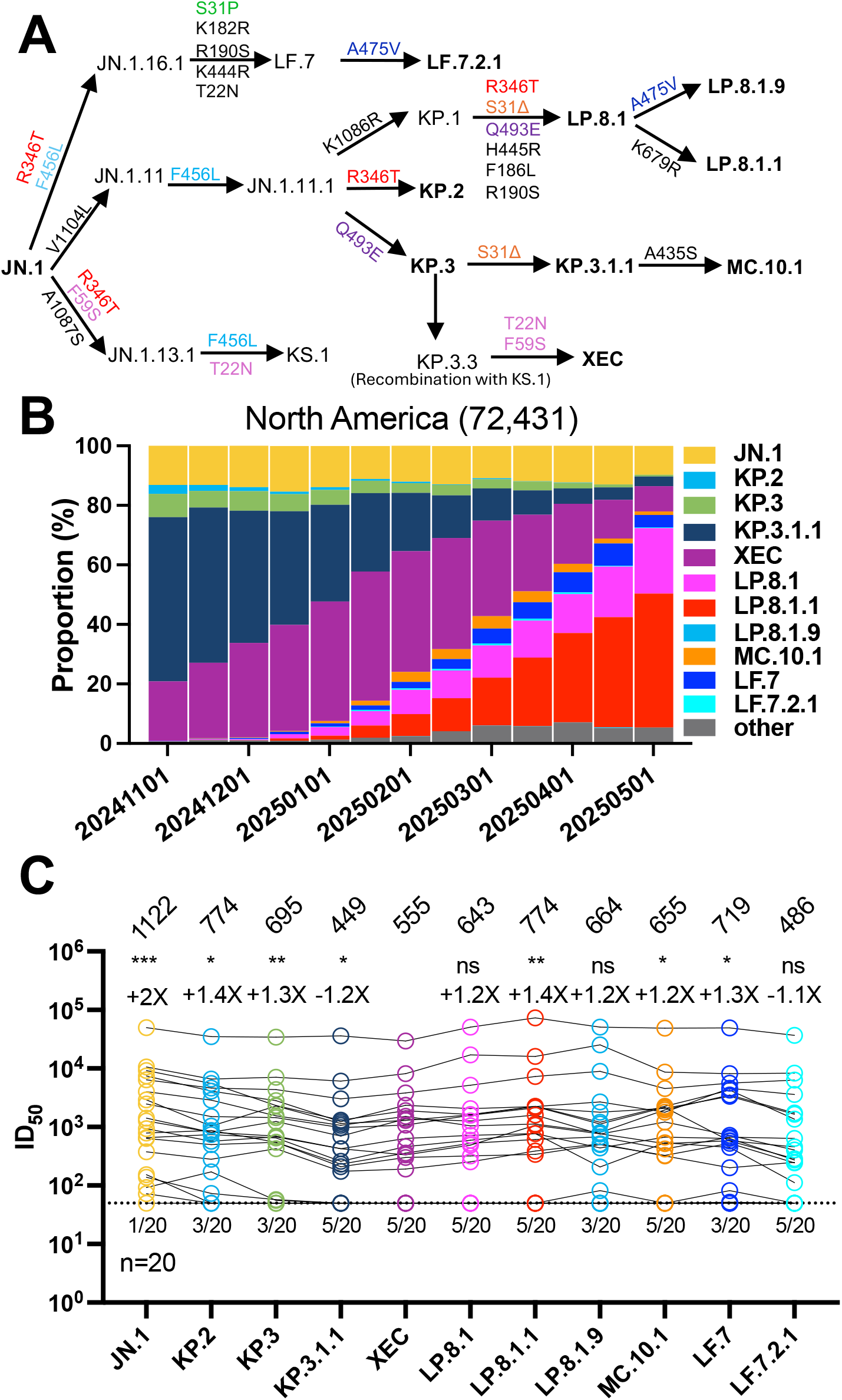
Evolution and serum neutralization of recently dominant SARS-CoV-2 JN.1 subvariants. **A**: Spike mutations present in recently dominant JN.1 subvariants. **B**: Relative frequencies of SARS-CoV-2 variants in North America from 11/01/2024 to 05/01/2025. Data from GISAID. **C**: Serum neutralizing titers (ID_50_) against VSV-based pseudoviruses bearing spike proteins of JN.1, KP.2, KP.3, KP.3.1.1, XEC, LP.8.1, LP.8.1.1, LP.8.1.9, MC.10.1, LF.7, or LF.7.2.1, for samples from recipients of KP.2 MV boosters at ∼1 month post-booster. The geometric mean ID_50_ titer (GMT) is presented at the top. The fold change in GMT for each virus compared to XEC is also shown immediately above the symbols. Statistical analyses used Wilcoxon matched-pairs signed-rank tests, comparing to XEC. n, sample size; ns, not significant. * p < 0.05, ** p < 0.01, *** p < 0.001, **** p < 0.0001. Numbers under the dotted lines denote numbers of serum samples that were under the limit of detection (ID_50_ < 50).

Another JN.1 progeny, LF.7, emerged in Asia, featuring 7 additional spike mutations beyond JN.1, including T22N, S31P, K182R, R190S, R346T, K444R, and F456L. LF.7.2.1, a descendant of LF.7 bearing a further A475V mutation, quickly outcompeted LF.7. However, LF.7.2.1 is not as dominant in North America (Fig. 1B). Another variant under monitoring, MC.10.1, includes A435S in addition to the other spike mutations found in KP.3.1.1 (Fig. 1A). MC.10.1 was gradually increasing in early 2025 but is no longer growing in any location (Figs. 1A, 1B). Due to the many spike mutations and recent dominance of these variants, an understanding of their antigenic properties, particularly of LP.8.1.1, is of great importance to regulatory agencies and vaccine advisory bodies as they consider whether COVID-19 vaccines need to be updated to provide better coverage. To assess the antigenicity of LP.8.1.1 and other recent subvariants, we performed pseudovirus neutralization assays using serum samples from 20 KP.2 MV recipients in the US obtained at approximately 1 month post boost (Tables S1, S2). We found that neutralizing titers against LP.8.1, LP.8.1.1, MC.10.1, LF.7, and LF.7.2.1 (geometric means 643, 774, 655, 719, 486, respectively) were at least equivalently high to those against XEC (geometric mean 555) in the serum of 20 US-based participants who received KP.2 MV (Fig. 1C). In a subset of these participants, we then checked for durability of neutralizing antibody titers. Since LP.8.1.1 is antigenically similar to LP.8.1 in the serum of KP.2 MV recipients, we focused on titers against LP.8.1. Serum neutralizing antibody titers against KP.3.1.1, XEC, and LP.8.1 were similar in range as well as in their respective geometric mean titers at 1 month post boost and at 4 months post boost (Fig. S1C). The estimated half-lives for the decay in neutralizing antibodies titers against these subvariants were also comparable, ranging from 66 to 91 days (Fig. S2). These findings show that LP.8.1, LP.8.1.1, MC.10.1, LF.7, and LF.7.2.1 are antigenically similar to KP.3.1.1 and XEC after KP.2 MV. Our finding of antigenic similarity of LP.8.1 is concordant with other published results using sera obtained from China, Japan, and Germany, in populations with different exposure and vaccination histories than those in North America ^3–5^. We also show that KP.2 MV recipients have titers against LP.8.1.1 that are at least as high as against LP.8.1. This conclusion, in turn, raises the specter that current COVID-19 boosters may not currently need reformulation. But antigenicity does not equal immunogenicity. Vaccine manufacturers could readily compare the immunogenicity of LP.8.1-based vaccines to that of KP.2 MV in both animals and humans. Comparable results from such studies would reinforce the suggestion provided herein that vaccine manufacturers could potentially avoid the production of a new formulation of COVID-19 booster shots for 2025, barring the emergence of another distinct SARS-CoV-2 variant in the meantime.

## Notes

### Competing Interest Statement

D.D.H. co-founded TaiMed Biologics and RenBio, and he serves as a consultant for WuXi Biologics and Brii Biosciences and is a board director at Vicarious Surgical. A.G. served as a member of the scientific advisory board for Janssen Pharmaceuticals and has consulted and serves on a scientific advisory board for Sanofi Pasteur. The remaining authors declare no conflicts of interest.

### Summary of Updates

Additional pseudovirus neutralization assays against a broader set of currently dominant or concerning SARS-CoV-2 viral variants, using samples from an enlarged cohort. Prevalence and structural diagram panels also updated accordingly. The overall conclusion remains concordant with the first version of the preprint: circulating variants are antigenically similar in the serum of KP.2-directed vaccine recipients, and therefore it may be acceptable to use KP.2-directed vaccine boosters in 2025, as well.

### Funding

- National Institutes of Health: subcontract no. 0258-A700-4609 under federal contract no. 75N93021C00014, subcontract GR0010139-PO024016 under federal contract no. 75N93021C00016, K08 AI180347, K23 AI171263, K24 AI155230
- Bill & Melinda Gates Foundation: INV019355
- Columbia University: UR014016

