## Supplementary Appendix for "Do Existing COVID-19 Vaccines Need to Be Updated in 2025?"

### Supplementary Methods

*Clinical cohorts*

Serum samples were collected through the VIVA study at the University of Michigan and through the "COVID-19 Persistence and Immunology Cohort (C-PIC)" study at Columbia University. Specimens were obtained following participant informed consent and in adherence to the protocols approved by the IRBs of University of Michigan Medical School (protocol HUM00232359) and Columbia University (protocol AAAS9722).

In this study, serum samples were collected from individuals who had been administered the KP.2 monovalent vaccine booster (KP.2 MV). Serum was collected at approximately 1 month and approximately 4 months after receiving the booster. The majority of the study subjects were female, representing 68.8%, with an average age of 39.5 years. Serum samples were collected, on average, 32.7 days and 112.7 days post KP.2 MV booster. Further demographic details, vaccination status, and serum collection timelines are summarized in Tables S1 and S2. NP ELISAs were performed, as previously described[1], to check for evidence of unreported infections between the two samples for each participant.

*Cell lines*

HEK293T (ATCC, CRL-3216) cells and Vero-E6 cells (ATCC, CRL-1586) were cultured in Dulbecco's Modified Eagle's Medium (DMEM; Sigma) that contained 10% heat-inactivated fetal bovine serum (FBS; Fisher) and 1% penicillin-streptomycin (PS; Fisher). All cell lines were cultured in an atmosphere of 5% $CO_2$ at 37°C.

*SARS-CoV-2 spike plasmids*

The spike constructs for JN.1, KP.3.1.1 and XEC were generated as previously reported[2–4]. We performed site-directed mutagenesis with Agilent QuikChange II XL and Agilent QuikChange Multi mutagenesis kits to generate the spike construct for LP.8.1, LP.8.1.1, MC.10.1, LF.7, and LF.7.2.1. All sequences were confirmed by whole-plasmid sequencing.

*Pseudotyped SARS-CoV-2 variants*

Pseudotyped SARS-CoV-2 was produced as previously described[5]. HEK293T cells were first transfected with spike-encoding plasmids using 1 mg/mL of PEI-MAX (Polysciences, Inc.) and cultured for 24 hours. The transfected HEK293T cells were then infected with VSV-G pseudotyped ΔG-luciferase virus (Kerafast, EH1020-PM) at a multiplicity of infection (MOI) of approximately 3 to 5. Two hours later, the cells were washed three times with and cultured in fresh competent cell medium overnight. The virus was then harvested first by centrifugation at 2000 rpm for 10 minutes followed by collection of the supernatant. 20% of I1 hybridoma (ATCC, CRL-2700) supernatant was then added to each virus before storing at -80°C until use.

*Pseudovirus serum neutralization and hACE2 inhibition*

Pseudoviruses were titrated to standardize the viral input prior to each neutralization assay. For neutralization assays, serum samples were inactivated at 56°C for 30 minutes before use, and the inactivated sera were serially diluted, from a starting dilution of 1:50, with a dilution factor of four, across 7 total serial dilutions. For ACE2 inhibition assays, as previously reported, soluble chimeric human ACE2 (hACE2), which contains ACE2 residues 1-732 and is fused to human IgG1 Fc, was diluted from 10 μg/mL with a dilution factor of three across 7 total serial dilutions. Following this, pseudoviruses were added and incubated at 37 °C for 1 hour. As a control, wells containing only the pseudovirus were also prepared on each test plate. Vero-E6 cells were then seeded at 40,000 cells per well and were incubated overnight at 37°C, on black Costar 96-well plates. Promega Luciferase Assay System (E4550) was used for lysis and luciferase activity measurements on a Tecan Infinite® 200 PRO using i-control™ software v.3.9.1.0, in accordance with the manufacturer's instructions. The serum dilution or hACE2 concentration that inhibits 50% of virus entry ($ID_{50}$ or $IC_{50}$, respectively) was calculated using five-parameter log-logistic dose-response curve fitting with the drda package (v2.0.5) in R[6].

*Quantification and statistical analysis*

$ID_{50}$ values at or below the serum neutralization assay dilution factor limit of detection (LOD) of 50 were treated as 50 for the purpose of calculating each group's geometric mean $ID_{50}$ titer (GMT). Half-life was estimated assuming exponential decay between the sample time points; half-life could not be estimated for participants with unchanged titers (both samples below LOD) or with apparent increase in titers. Statistical significance of differences between viruses was evaluated using the paired Wilcoxon signed-rank test in R (v4.3.2). Some plots were subsequently generated in GraphPad Prism for clarity. Levels of significance are denoted on figures as follows: ns, not significant; *$p < 0.05$, **$p < 0.01$, ***$p < 0.001$, and ****$p < 0.0001$.

### Acknowledgements

This study was supported by funding from the NIH SARS-CoV-2 Assessment of Viral Evolution (SAVE) Program (subcontract no. 0258-A700-4609 under federal contract no. 75N93021C00014 to D.D.H. and (subcontract GR0010139-PO024016 under federal contract no. 75N93021C00016) to A.G. and the Gates Foundation (project INV019355) to D.D.H., internal startup funding UR014016 from Columbia University to Y.G. K08 AI180347 to A.B., K23 AI171263 to L.J.P., K24 AI155230 to M.T.Y. We thank all who contributed their data to the Global Initiative on Sharing All Influenza Data (GISAID).

We thank Amanda Castillo, Meredith McNairy and Antonia Sturiza for conducting the C-PIC study (Columbia), and to Zijin Chu, Theresa Kowalski-Dobson, Anna Buswinka, Gabe Simjanovski, Joseph Wendzinski, Mayurika Patel, Kathleen Lindsey, and Dawson Davis of the VIVA study team for conducting the VIVA study (at University of Michigan).

### Author Contributions

The study was conceptualized by I.A.M., Y.G., Q.W., and D.D.H. Experiments were conducted and data analyzed by I.A.M., M.W., and Y.G. Project management was handled by I.A.M. Serum samples were collected by I.A.M., A.B., C.G., V.M.P., J.G.S., L.J.P., M.T.Y., A.G., and their colleagues. The results were analyzed, and the manuscript was written by I.A.M., M.W., Y.G., and D.D.H. All contributing authors have reviewed and endorsed the manuscript.

### Declaration of Interests

D.D.H. co-founded TaiMed Biologics and RenBio, and he serves as a consultant for WuXi Biologics and Brii Biosciences and is a board director at Vicarious Surgical. A.G. served as a member of the scientific advisory board for Janssen Pharmaceuticals and has consulted and serves on a scientific advisory board for Sanofi Pasteur. The remaining authors declare no conflicts of interest.

### Supplementary Figures

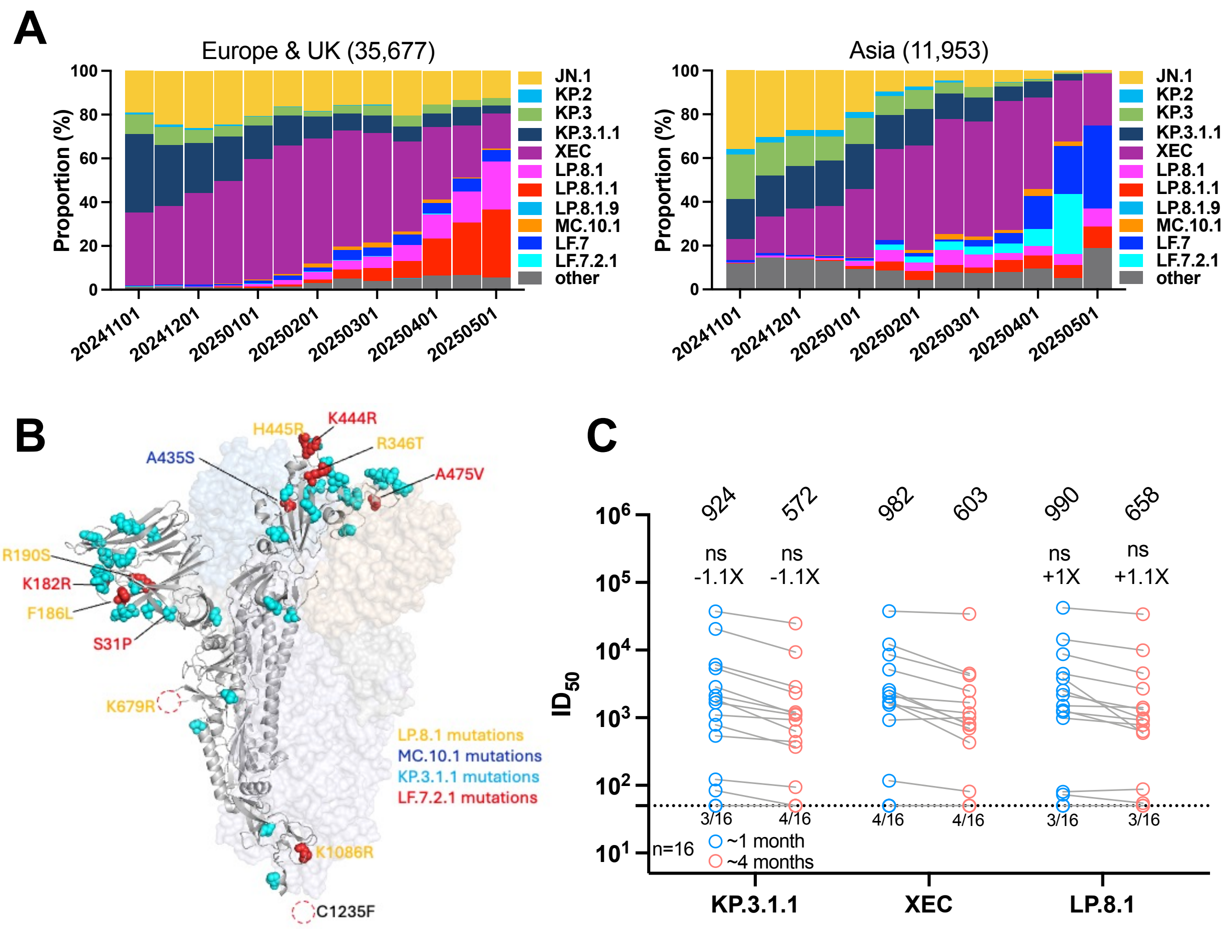

*Figure S1: Frequencies of SARS-CoV-2 variants, structural diagram of related mutations, and titers against LP.8.1 over time*. **A**: Relative frequencies of SARS-CoV-2 variants from 11/01/2024 to 05/01/2025 in the indicated regions. Data from GISAID. **B**: Structural diagram of SARS-CoV-2 spike protein. Mutations present in KP.3.1.1 highlighted in cyan. Additional mutations present in LP.8.1 highlighted in yellow. K679R mutation in LP.8.1.1 highlighted in yellow text with dashed circle. Additional mutation present in MC.10.1 highlighted in blue. Mutations present in LF.7.2.1 highlighted in red. **C**: Serum neutralizing titers ($ID_{50}$) against VSV-based pseudoviruses bearing spike proteins of KP.3.1.1, XEC, or LP.8.1, for samples from recipients of KP.2 MV boosters at ~1 month and ~4 months time points post-booster. The geometric mean $ID_{50}$ titer (GMT) is presented at the top. The fold change in GMT for each virus compared to XEC is also shown immediately above the symbols. Statistical analyses used Wilcoxon matched-pairs signed-rank tests, comparing to XEC. n, sample size; ns, not significant. * $p < 0.05$, ** $p < 0.01$, *** $p < 0.001$, **** $p < 0.0001$. Numbers under the dotted lines denote numbers of serum samples that were under the limit of detection ($ID_{50} < 50$).

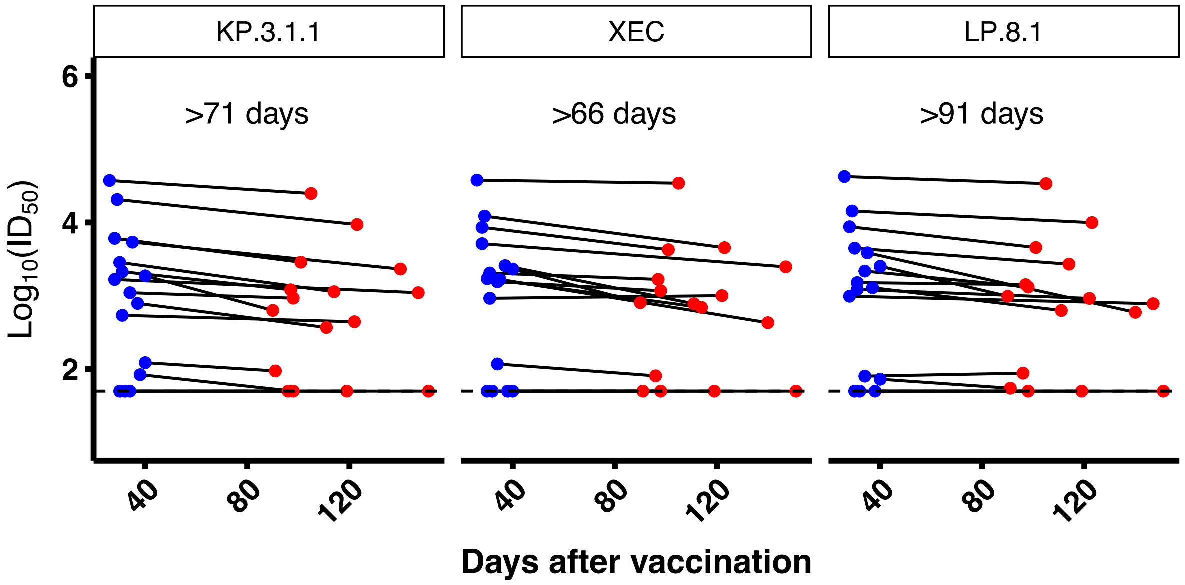

***Figure S2**: Serum neutralizing titers over time and estimated half-life*. Paired $ID_{50}$ values per participant. Geometric mean estimated half-life of titers noted for each variant. Half-life was not estimated for participants with unchanged $ID_{50}$ (i.e., both values below LOD), nor for the one participant who had an apparent small increase in titer. Blue = first sampling post-boost, Red = second sampling post-boost.

### Supplementary Tables

| ID | Group | Age (Yr) | Sex | Race | No. Vax | No. WT Vax | No. BA.5 Bivalent Vax | No. XBB.1.5 Vax | No. KP.2 MV | Sera Days Post Most Recent Infx | Sera Days Post Most Recent Vaccine | Vaccine History |
|---|---|---|---|---|---|---|---|---|---|---|---|---|
| * CUMC 1 | KP.2 MV | 25 | F | As | 6 | 3 | 1 | 1 | 1 | 880 | 31 | WT-P/WT-P/WT-P/BA.5-M/XBB.1.5-M/KP.2-M |
| * CUMC 2 | KP.2 MV | 35 | M | As | 4 | 3 | 0 | 0 | 1 | - | 26 | WT-P/WT-P/WT-P/KP.2-M |
| * CUMC 3 | KP.2 MV | 21 | M | As | 4 | 3 | 0 | 0 | 1 | 195 | 37 | WT-P/WT-P/WT-P/KP.2-P |
| * CUMC 4 | KP.2 MV | 29 | F | As | 4 | 3 | 0 | 0 | 1 | 244 | 28 | WT-P/WT-P/WT-P/KP.2-M |
| * CUMC 5 | KP.2 MV | 55 | M | As | 5 | 3 | 1 | 0 | 1 | 451 | 30 | WT-M/WT-M/WT-M/BA.5-P/XBB.1.5-P/KP.2-P |
| * CUMC 6 | KP.2 MV | 36 | M | Wh | 6 | 3 | 1 | 1 | 1 | 796 | 30 | WT-P/WT-P/WT-P/WT-P/BA.5-P/XBB.1.5-P/KP.2-M |
| * CUMC 8 | KP.2 MV | 23 | F | As | 4 | 3 | 0 | 0 | 1 | - | 40 | WT-P/WT-P/WT-P/KP.2-M |
| * CUMC 9 | KP.2 MV | 42 | M | As | 6 | 3 | 1 | 1 | 1 | 650 | 34 | WT-P/WT-P/BA.5-P/XBB.1.5-P/KP.2-M |
| * CUMC 11 | KP.2 MV | 61 | F | Wh | 5 | 3 | 1 | 0 | 1 | 826 | 34 | WT-P/WT-P/WT-P/BA.5-P/KP.2-P |
| * UMICH 1 | KP.2 MV | 29 | F | Af | 6 | 3 | 1 | 1 | 1 | 784 | 31 | WT-P/WT-P/WT-P/BA.5-P/XBB.1.5-M/KP.2-P |
| * UMICH 2 | KP.2 MV | 24 | F | Wh | 6 | 3 | 1 | 1 | 1 | 658 | 28 | WT-P/WT-P/WT-P/BA.5-P/XBB.1.5-M/KP.2-P |
| * UMICH 3 | KP.2 MV | 25 | F | Wh | 6 | 3 | 1 | 1 | 1 | - | 32 | WT-P/WT-P/WT-M/BA-M/XBB.1.5-M/KP.2-P |
| * UMICH 4 | KP.2 MV | 80 | F | Wh | 8 | 4 | 1 | 2 | 1 | - | 40 | WT-P/WT-P/WT-P/WT-P/BA.5-P/XBB.1.5-P/XBB.1.5-P/KP.2-P |
| * UMICH 5 | KP.2 MV | 33 | F | As | 6 | 3 | 1 | 1 | 1 | - | 38 | WT-P/WT-P/WT-P/BA.5-M/XBB.1.5-P/KP.2-P |
| * UMICH 6 | KP.2 MV | 55 | F | Wh | 6 | 3 | 1 | 1 | 1 | 653 | 29 | WT-M/WT-M/WT-M/BA.5-M/XBB.1.5-M/KP.2-M |
| * UMICH 7 | KP.2 MV | 59 | F | Wh | 7 | 4 | 1 | 1 | 1 | - | 35 | WT-P/WT-P/WT-P/WT-P/BA.5-P/XBB.1.5-M/KP.2-P |
| UMICH 8 | KP.2 MV | 64 | M | Wh | 6 | 3 | 1 | 1 | 1 | 741 | 31 | WT-P/WT-P/WT-P/BA.5-P/XBB.1.5-P/KP.2-P |
| UMICH 9 | KP.2 MV | 66 | F | Wh | 6 | 3 | 1 | 1 | 1 | 1685 | 31 | WT-M/WT-M/WT-P/BA.5-P/XBB.1.5-P/KP.2-P |
| UMICH 10 | KP.2 MV | 27 | F | Wh | 5 | 2 | 1 | 1 | 1 | 476 | 37 | WT-J/WT-M/BA.5-M/XBB.1.5-M/KP.2-M |
| UMICH 11 | KP.2 MV | 61 | F | Wh | 7 | 4 | 1 | 1 | 1 | 380 | 50 | WT-P/WT-P/WT-P/WT-P/BA.5-P/XBB.1.5-P/KP.2-M |

*Table S1: Individual participant characteristics and sample timing*. Vaccine formulations are denoted as wildtype (WT), BA.5 Bivalent (BA.5), XBB.1.5 monovalent (XBB.1.5), and KP.2 monovalent (KP.2). Vaccine manufacturers are denoted as Pfizer (P), Moderna (M). As, Asian; Wh, White; Af, Black or African American; Yr, years; Infx, infection; Vax, vaccination. * in ID field indicates that the ~1-month and ~4-month sample was used in Figs. S1C and S2, in addition to in other figures.

| | | All Participants | |
|---|---|---|---|
| | | No. or Mean | % or (range) |
| **Total** | | 20 | - |
| **Female** | | 14 | 70.0% |
| **Male** | | 6 | 30.0% |
| **Age** | | 42.5 | (21, 80) |
| | All vaccines | 5.7 | (4, 8) |
| | WT | 3.1 | (2, 4) |
| | BA.5 BV | 0.8 | (0, 1) |
| | XBB.1.5 | 0.8 | (0, 2) |
| **No. Vaccines** | KP. 2 MV | 1.0 | (1,1) |
| **Sera Days Post Most Recent Infection** | | 672.8 | (0, 1685) |
| **Sera Days Post Most Recent Vaccination** | | 33.6 | (26, 1208) |

*Table S2: Summary of the clinical cohort*. Vaccine formulations are denoted as wildtype (WT), BA.5 Bivalent (BA.5), XBB.1.5 monovalent (XBB.1.5), and KP.2 monovalent (KP.2).
